# *Cis*-regulatory evolution in prokaryotes revealed by interspecific archaeal hybrids

**DOI:** 10.1101/044362

**Authors:** Carlo G. Artieri, Adit Naor, Israela Turgeman-Grott, Yiqi Zhou, Ryan York, Uri Gophna, Hunter B. Fraser

## Abstract

The study of allele-specific expression (ASE) in interspecific hybrids has played a central role in our understanding of a wide range of phenomena, including genomic imprinting, X-chromosome inactivation, and cis-regulatory evolution. However across the hundreds of studies of hybrid ASE, all have been restricted to sexually reproducing eukaryotes, leaving a major gap in our understanding of the genomic patterns of cis-regulatory evolution in prokaryotes. Here we introduce a method to generate stable hybrids between two species of halophilic archaea, and measure genome-wide ASE in these hybrids with RNA-seq. We found that over half of all genes have significant ASE, and that genes encoding kinases show evidence of lineage-specific selection on their cis-regulation. This pattern of polygenic selection suggested species-specific adaptation to low phosphate conditions, which we confirmed with growth experiments. Altogether, our work extends the study of ASE to archaea, and suggests that cis-regulation can evolve under polygenic lineage-specific selection in prokaryotes.

## Introduction

For the past 50 years, interspecific hybrids have been an invaluable resource for studying the regulation of gene expression. Beginning with studies in species such as frogs and trout, the concept of allele-specific expression (ASE) was first used to investigate differences in enzyme activity levels for the two alleles in a hybrid (Wright and Moyer 1966, Hitzeroth et al. 1968). Since then, measurement of ASE in hybrids has played a critical role in the study of genomic imprinting, X-chromosome inactivation, and cis-regulatory evolution (Avise and Duvall 1977, Dickinson et al. 1984, Bartolomei et al. 1991, Wittkopp et al. 2004, Wang et al 2010, Pastinen 2010, Wittkopp and Kalay 2011, Wang and Clark 2014, Babak et al 2015).

Particularly since the advent of high-throughput RNA-sequencing (RNA-seq), ASE in hybrids has been a major focus for studies of gene expression evolution. In a hybrid, the two alleles of each gene are present in the same cells, and thus experience all the same environmental factors/perturbations, which makes direct comparison more meaningful than when the expression profiles of different species are compared—especially when the environments of those species are not well-controlled, such as in human studies. In addition, because the two alleles in a hybrid are exposed to all the same trans-acting factors (such as transcription factors)—which can affect gene expression levels, but cannot cause allelic bias in the absence of cis-regulatory divergence—ASE reflects only cis-acting differences between alleles, which can be a useful distinction for a wide range of questions in evolutionary biology (Wittkopp and Kalay 2011).

Despite the multitude of studies employing ASE (over 650 publications when searching “allele-specific expression” or “allele-specific gene expression” in PubMed abstracts), a limitation shared by all of them is that they have been restricted to eukaryotes. The reason for this is that prokaryotes do not undergo sexual reproduction, so generating hybrids has not been possible. As a result, our knowledge of cis-regulatory evolution in prokaryotes has lagged far behind that in eukaryotes.

However, some halophilic archaea can undergo a fusion process that can generate hybrid cells (Rosenshine et al. 1989, Ortenberg et al. 1999). This process is efficient even between different species, but the heterozygous hybrid state is unstable due to gene conversion events (Lange et al. 2011), as well as large-scale recombination events that result in homozygous recombinants (Naor et al. 2012). We overcame this obstacle by maintaining two *different* selection markers at the same genetic locus in the two parental species. In such a condition any homologous recombination event will result in swapping one selection marker for the other, and as long as one selects for *both* markers, only heterozygous cells will be able to keep dividing, assuming no ectopic recombination occurs.

We have applied this unique system to explore cis-regulatory evolution in the genus *Haloferax.* The two species we studied were *Haloferax volcanii*, isolated from the Dead Sea in Jordan (Mullakhanbhai and Larsen 1975), and *Haloferax mediterranei*, isolated from a saltern in Alicante, Spain (Rodriguez-Valera et al. 1983). While both isolation sites were characterized by high salt concentrations, they likely differed greatly in other respects, such as concentrations of mangesium and phosphate ions, raising the possibility of lineage-specific adaptations of these species to their respective environments.

## Results

We have previously shown that *H. volcanii* and *H. mediterranei* are able to efficiently mate and generate interspecies recombinants (Naor et al. 2012). In order to generate a stable *H. volcanii* - *H. mediterranei* hybrid, we needed to prevent the possibility of recombination between chromosomes, thus forcing the hybrid to retain both parental chromosomes. For that we needed to create mutants that carry two different selectable markers at the same genomic location, since the two strains are syntenic (López-García et al. 1995) (Figure 1A). We used the *H. mediterranei* strain WR646 (*ΔtrpA hdrB*+), an auxotroph for tryptophan and prototroph for thymidine (Naor et al 2012), and the *H. volcanii* strain H133 (*ΔtrpA ΔhdrB*), an auxotroph for tryptophan and thymidine (Allers et al. 2004). H133 was then modified by inserting the *trpA* selectable marker into the *hdrB* locus to generate UG241 (*trpA*+ *ΔhdrB*). This was done by transforming H133 with pTA160-trpA and selecting on media lacking thymidine, thus selecting for a double crossover event copying the *trpA* selectable marker into the *hdrB* locus. To create the stable hybrid, WR646 and UG241 were mated and colonies were selected on media lacking thymidine and tryptophan (Figure 1B and Methods).

**Figure 1.**
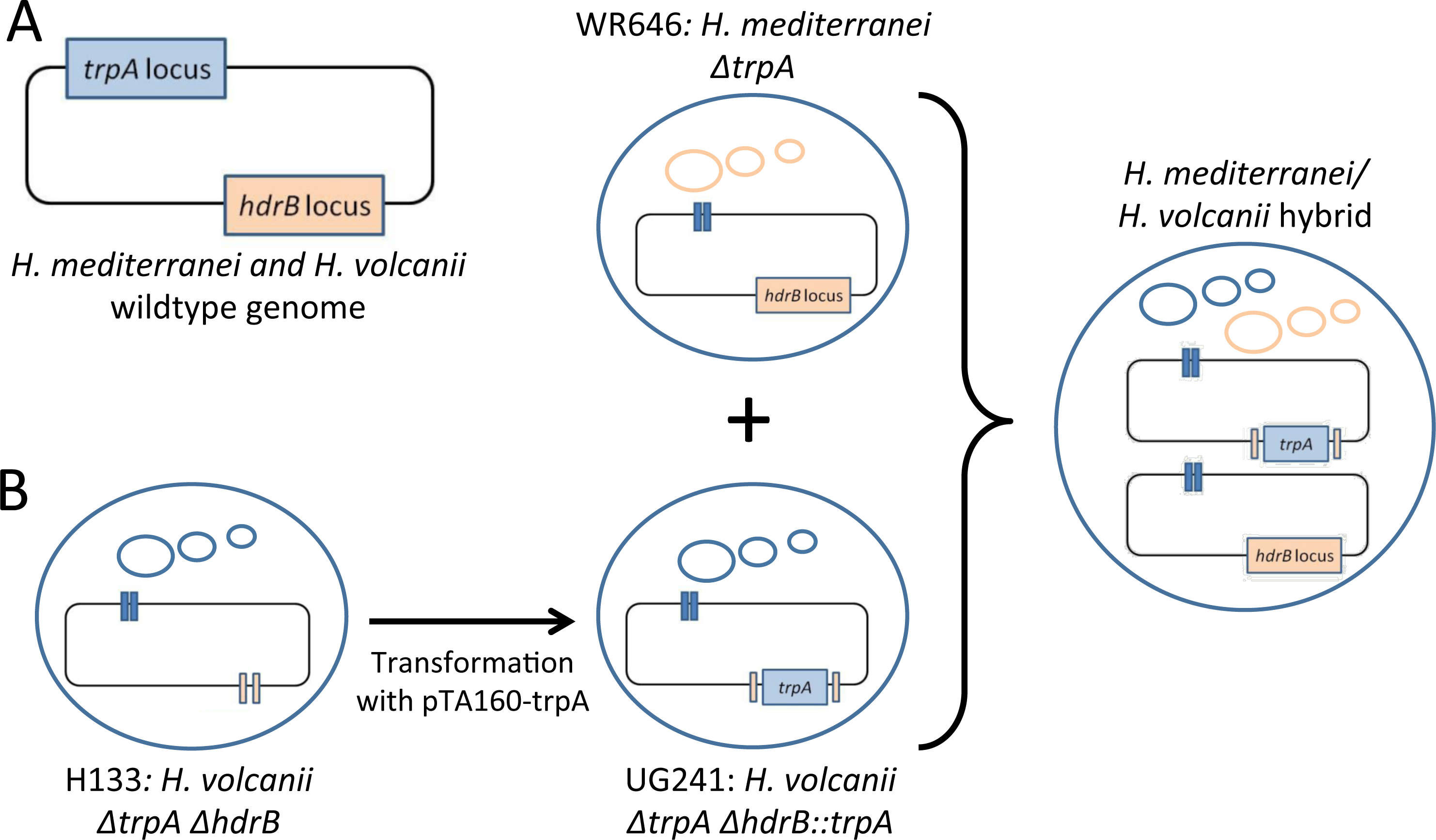
Generation of stable *H. volcanii* - *H. mediterranei* hybrids. **A.** The genomic organization of the selectable markers involved in the study. **B.** Generation of a stable hybrid. H133 was transformed with pTA160 *trpA*, and upon selection on media lacking thymidine the *trpA* marker was integrated in the *hdrB* locus, generating UG241. UG241 was mated with WR646, which are autotrophs for thymidine and tryptophan, respectively. The mated colonies were selected on a media lacking thymidine and tryptophan. Small circles indicate the plasmids and the rectangle represents the chromosome.

We performed both RNA- and DNA-seq on two independently derived *H. volcanii* × *H. mediterranei* hybrid cultures (hereafter replicates 1 and 2). Reads were mapped to a reference containing both parental genomes, and species-specific gene-level expression was calculated in <u>r</u>eads <u>p</u>er <u>k</u>ilobase per <u>m</u>illion mapped reads (RPKM). Integrating ortholog and operon predictions (Price et al. 2005; DeLuca et al. 2012), resulted in 1,954 orthologous transcriptional units, hereafter referred to as ‘orthologs’, corresponding to 1,507 individual genes and 447 operons (Supplemental File 1; see Methods).

As *Haloferax* species are highly tolerant of both intra- and inter-chromosomal and plasmid copy number variation (Breuert et al. 2006), we used the DNA-seq data to identify large-scale amplifications. As expected, the ratio of plasmid coverage to chromosomal coverage varied among species and between replicates (Supplemental Fig. 1). Consequently, we restricted our analysis to orthologs found outside of amplified regions and on the main chromosomes of the two parental species, resulting in 1,526 orthologs for analysis (Supplemental Table 1). We observed similar patterns of expression levels and ASE ratios in the two biological replicates (Supplemental Fig 2).

Differential expression of the two species’ alleles within a common *trans* cellular background, known as allele-specific expression (ASE), indicates divergence of *cis*-regulation between orthologs (Pastinen 2010, Wittkopp and Kalay 2011). In order to detect significant ASE, we employed a method that takes into account both gene length and base-compositional differences between parental alleles (see Methods) (Bullard et al. 2010; Artieri and Fraser 2014). 929 orthologs showed significant ASE at a 5% false-discovery rate (FDR), indicating the presence of substantial cis-regulatory differences between the two parental species (Figure 2A). We found no significant difference in the number of genes favoring either species’ allele (453 vs. 476 favoring the *H. mediteranei* vs. *H. volcanii* allele, χ^2^ = 0.569, 1 degree of freedom, p = 0.451), suggesting that ASE was about equally likely to favor either allele.

**Figure 2.**
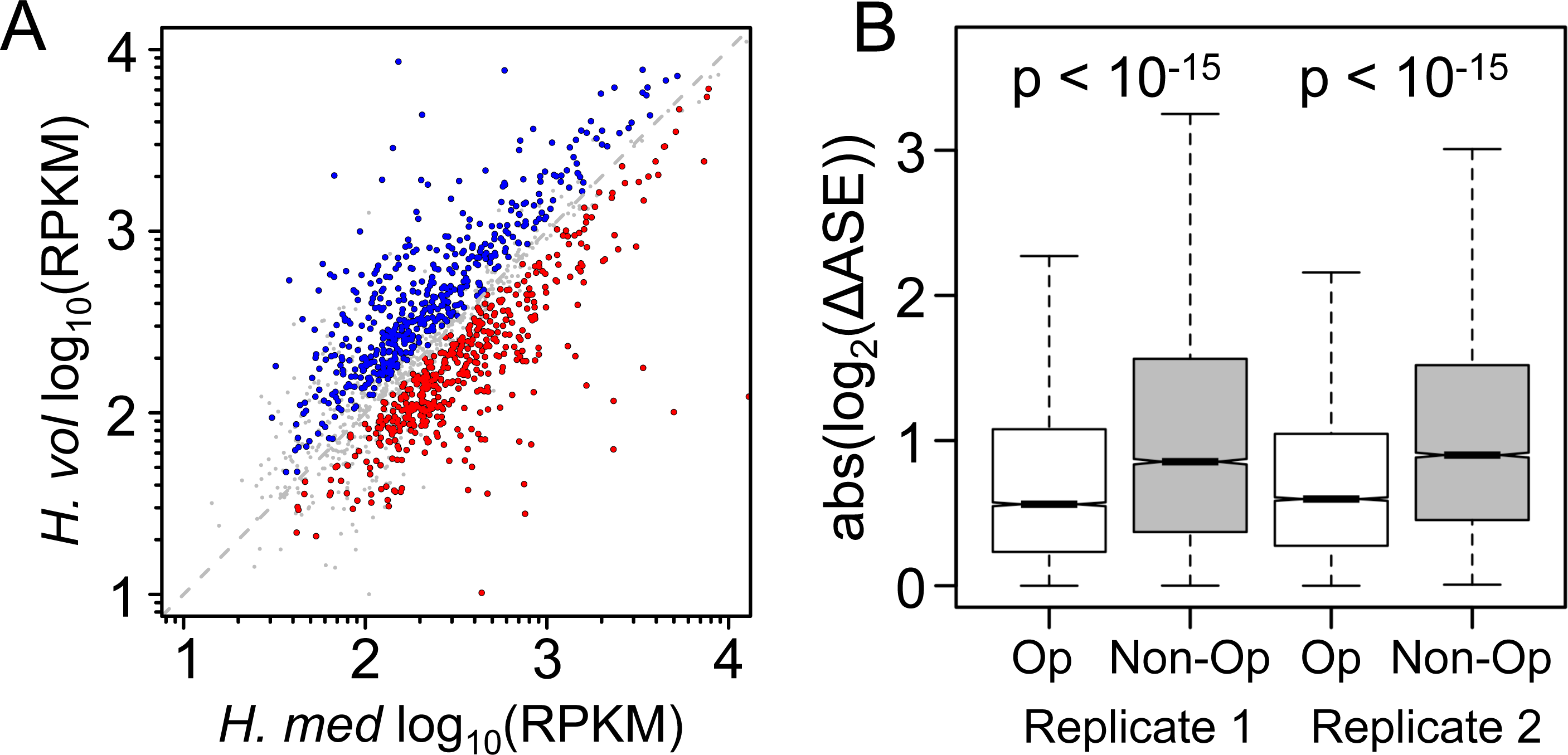
Regulatory divergence between archaeal hybrids is revealed by ASE analysis. **(A)** Approximately equal numbers of orthologs show significant allelic bias favoring either the *H. mediterranei* (453, red) or *H. volcanii* allele (476, blue). RPKMs plotted in this figure are the mean of the two biological replicates after normalization (see Methods). med, mediterranei; vol, volcanii. **(B)** Pairwise comparisons of adjacent genes within predicted operons show significantly more similar ASE than independently transcribed adjacent genes (Kruskal-Wallis rank sum test, p < 10^-15^ in both cases). Op, adjacent genes within predicted operons; Non-Op, adjacent genes outside of predicted operons.

We also tested the accuracy of our classification of genes into orthologous operons by asking whether adjacent genes within operons showed greater similarity in ASE ratios than adjacent, independently transcribed genes. Indeed, genes within operons had a significantly smaller median absolute log_2_ differences in ASE values than those outside of operons in both biological replicates (Figure 2B; Kruskal-Wallis test p < 10^-15^ for each replicate). These differences may be conservative, since any errors in the operon predictions (Price et al. 2005) would lead us to underestimate their magnitudes.

Although ASE data reveal genome-wide patterns of cis-regulatory divergence, these might mostly reflect random changes due to genetic drift of neutral alleles. To identify those changes driven by lineage-specific natural selection, we and others have developed a “sign test” that detects selection acting on the regulation of entire groups of functionally related genes (reviewed in Fraser 2011). This test has been successfully applied to fungi, plants, and metazoans (Fraser et al. 2010, Bullard et al. 2010, Fraser et al. 2011, Fraser et al. 2012, Fraser 2013, House et al. 2014, Naranjo et al. 2015), but not to prokaryotes, due to the previous lack of ASE data from interspecific hybrids.

We applied the sign test to Gene Ontology gene sets from *H. volcanii* (Binns et al. 2009), to search for gene sets with ASE directionality biased towards one parental species (see Methods). Such a directionality bias represents a robust signature of lineage-specific selection (Fraser 2011). We found that genes with a known role in phosphorylation (G0:0016310) showed a significant bias in ASE directionality (ASE for 16/21 alleles favoring *H. mediterranei* in each biological replicate; permutation-based p < 0.001). These phosphorylation-related genes were predominantly kinases, and the “kinase activity” subset (G0:0016301) showed a similar ASE bias (ASE for 15/19 alleles favoring *H. mediterranei* in each biological replicate; permutation-based p < 0.001; Figure 3A, Supplemental Table 1). This suggested that genes related to phosphorylation—particularly kinases—have evolved under lineage-specific selective pressures leading to increased expression in *H. mediterranei*, or decreased expression in *H. volcanii*.

**Figure 3.**
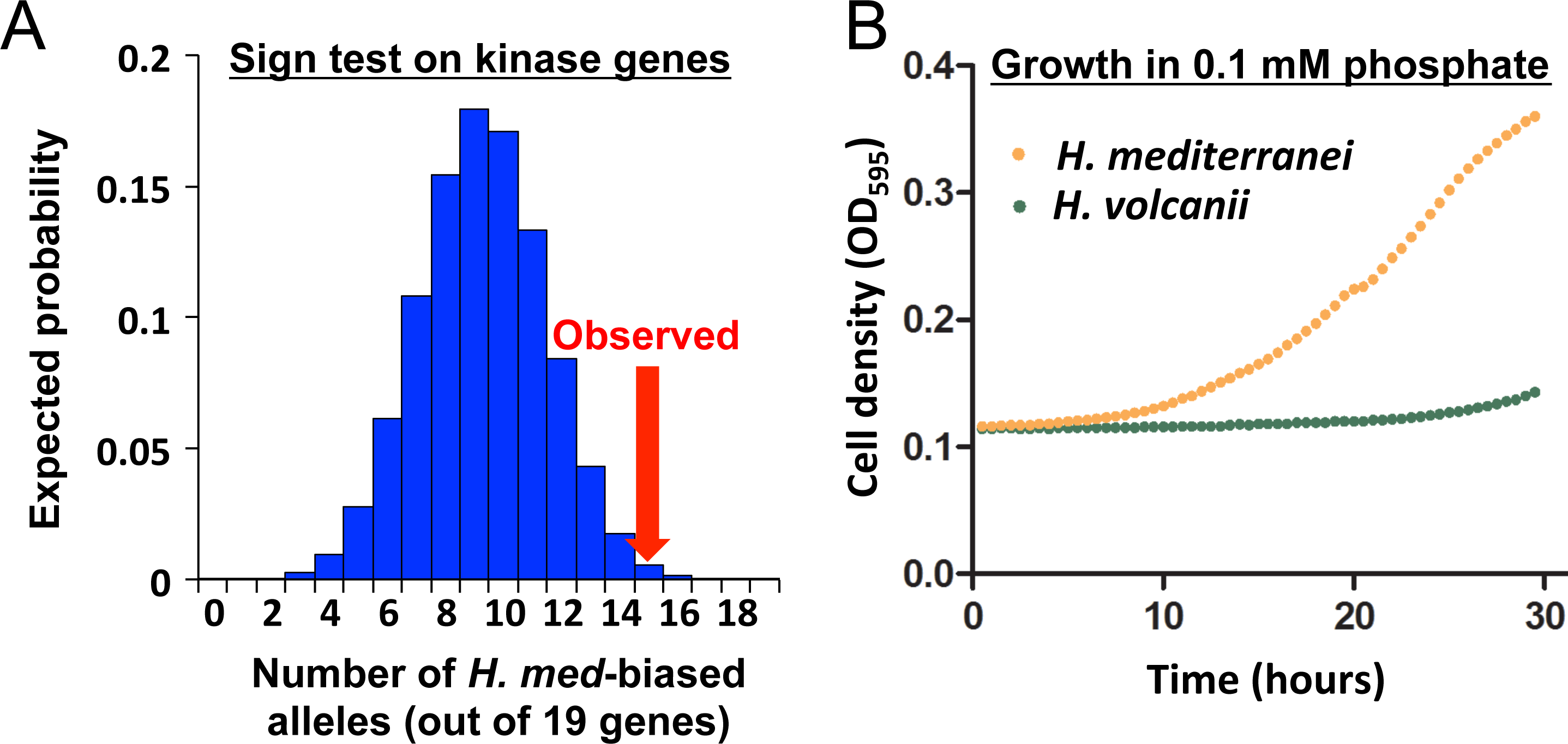
Detection of lineage-specific selection and differential fitness in low phosphate conditions. **(A)** For a set of 19 genes, the expected number with ASE with higher expression from the *H. mediterranei* alleles is plotted. The kinase gene set had 15/19 genes favoring the *H. mediterranei* alleles (red arrow), in both biological replicates. **(B)** *H. mediterranei* grows robustly in 0.1 mM phosphate, whereas *H. volcanii* does not. See also Supplemental Figure 3.

The results of the sign test led us to hypothesize that the higher expression of kinases in *H. mediterranei* may increase the fitness of this species in conditions where phosphate is limiting, since it could utilize the scarce phosphate more efficiently. To test this hypothesis, we grew both parental strains in low (0.1 mM) phosphate for 30 hours. Consistent with our prediction, *H. mediterranei* showed robust growth in this condition, in contrast to *H. volcanii*, whose growth was highly impaired (Figure 3B and Supplementary Figure 3).

## Conclusion

In this work we have introduced a method to create stable interspecific hybrids of *Haloferax*, and used these hybrids to investigate the extent and phenotypic impacts of cis-regulatory evolution. Our application of the sign test revealed lineage-specific selection acting on the cis-regulation of kinases, which led to our prediction—and confirmation—of *H. mediterranei*’s superior fitness in phosphate-limiting conditions.

Although we do not know the phosphate concentrations of the specific sites where these two species were isolated, it is well established that phosphate is the main limiting nutrient in the Mediterranean (Lazzari et al. 2016). In contrast, the Dead Sea contains higher phosphate levels, particularly in the sediments where *H. volcanii* was most abundant (Nissenbaum et al. 1990). Therefore it is not surprising that *H. mediterranei* may have adapted to the low phosphate levels with increased expression levels of kinases, compared to *H. volcanii*.

Over half (929/1526) of the orthologs we studied showed significant ASE, though this fraction would likely increase with greater sequencing depth. Based on this lower bound estimate, we conclude that cis-regulatory divergence is likely to be a major source of evolutionary novelty in *Haloferax*, and that polygenic selection can act on the regulation of groups of functionally related genes in prokaryotes, similar to patterns of polygenic adaptation that have been discovered with the sign test across a wide range of eukaryotes.

## Materials and Methods

### Generation of the hybrids

Strains used: WR646 (*ΔpyrE ΔtrpA hdrB*+), H133 (*ΔApyrE ΔtrpA ΔleuB ΔhdrB*), UG241 (*ΔpyrE ΔleuB ΔhdrB::trpA*). Plasmids used: pTA160 for *ΔhdrB* deletion in *ΔpyrE2* background (Allers et al. 2004), and pTA298 for making deletions in *ΔtrpA* background (Stroud et al. 2012).

Strains were routinely grown in rich medium (Hv-YPC). When selection was needed we used Casamino Acids medium (Hv-Ca). When required, 50 *μ*g/ml of thymidine, uracil or tryptophan were added. Following mating all strains were grown on enhanced Casamino broth. All media were made as described (http://www.haloarchaea.com/resources/halohandbook/ version 7.2). All growth was at 45°C unless otherwise noted.

To introduce *trpA* at the *hdrB* locus of *H. volcanii*, we first inserted the *trpA* gene into the plasmid pTA160, originally designed to delete *hdrB*. The *trpA* gene, under the ferredoxin (*fdx*) promoter of *H. salinarium*, was amplified using primers AP389 (aaagctagcgctcggtacccggggatcc) and AP390 (tttgctagccgttatgtgcgttccggat), from pTA298. Using NheI, the PCR product was inserted into pTA160 between the *hdrB* flanking regions. Transformation of *H. volcanii* was carried out using the PEG method as described in (Cline et al. 1989).

Prior to hybridization, each culture was grown to an OD_600_ of 1–1.1, and 2 ml samples were taken from both strains and applied to a 0.2 mm filter connected to a vacuum to eliminate excess media. The filter was then placed on a Petri dish containing a rich medium (HY medium + thymidine) for 48 hr at 42°C. The cells were washed and resuspended in Casamino broth, washed twice more in the same media, and plated on selective media.

### Sequence library construction

RNA was isolated using EZ-RNA Total RNA Isolation Kit (Biological Industries Cat.# 20-400). DNA purification was done using the spooling method as described (http://www.haloarchaea.com/resources/halohandbook/ version 7.2).

RNA-seq and DNA-seq libraries were prepared using Illumina TruSeq kits, following manufacturer protocols. All libraries were sequenced as multiplexed samples in one lane of an Illumina HiSeq 2000.

### Genome annotation and read mapping

We obtained the genome assemblies and annotations for *H. volcanii* (strain DS2) and *H. mediterranei* (strain ATCC 33500) from NCBI RefSeq (accession numbers: GCF_000025685.1 and GCF_000337295.1, respectively). In order to determine which bases in each genome would be unambiguously mappable in the hybrids, in each parental genome, we employed a sliding window of 75 bp (our mapping read length; see below) and a step of one bp to create simulated NGS reads. These reads were mapped to a reference consisting of both parent’s genomes using Bowtie 0.12.8, with default parameters, retaining only uniquely mapping reads. Any base overlapped by reads that could not be mapped uniquely were masked from further analysis (corresponding to 3.9% and 1.3% of the *H. volcanii* and *H. mediterranei* genomes, respectively.)

We identified orthologous genes between the two species using the RoundUp database (DeLuca et al. 2012). Genes were then grouped into operons based on the MicrobesOnline operon predictions in *H. volcanii* (Price et al. 2005; http://meta.microbesonline.org/operons/gnc309800.html). Corresponding *H. mediterranei* operons were inferred from the presence of co-linearity of orthologs between the parental species.

All DNA-seq and RNA-seq reads were trimmed to 75 bp in length and mapped to a reference consisting of the concatenation of both parental genomes using Bowtie, version 0.12.8, with default parameters and retaining only uniquely mapping reads. DNA-seq RPKM was calculated using the number of unambiguously mappable bases as the gene length.

### Detecting significant ASE

We determined base-level coverage of CDSs of both species for all uniquely mappable positions for both hybrid replicates for main chromosome located orthologs with at least 100 reads mapping among both alleles in both biological replicates (1,320 orthologs). As the DNA-seq data indicated that parental chromosomal abundance was not necessarily equal in both replicates, base-level coverages were linearly scaled such that the total coverage was identical for each species’ main chromosome. The RNA-seq RPKMs were calculated as the base level coverage / (the number of uniquely mappable bases × the total base level coverage for all orthologs × the mapped read length [75 bp]). We applied the resampling test of Bullard et al. (2010): the base-level read coverage of each parental allele was resampled with replacement 10,000 times, under two conditions: 1) using the *H. volcanii* marginal nucleotide frequencies (ð_v_ = ð_v_[A], ð_v_[C], ð_v_[G], ð_v_[T]) and the *H. volcanii* length, L_v_, and 2) using the *H. mediterranei* marginal nucleotide frequencies ð_m_ and the *H. mediterranei* length, L_m_. A started log_2_ ratio (total base level coverage from ð_v_,L_v_ + 1 / total base level coverage from ð_v_,L_v_ + 1), denoted as log_2_(^Hv+1^/_Hm+1_), was calculated from each resampling, and represents the variation in log_2_(^Hv+1^/_Hm+1_) cis-ratios expected between alleles due only to differential base frequencies and length. Both null distributions (one per allele) were compared against the observed started log_2_(^Hv+1^/_Hm+1_) cis-ratio in order to obtain a two-tailed p value based on how often the observed ratio was outside of the bounds of the null distribution. In cases where both replicates agreed in the direction of parental bias, the least significant p-value among the four comparisons (both alleles in each replicate) was retained as a measure of the significance of differential expression. All p-values were adjusted such that we retained only those comparisons significant at an FDR of 5% for further analysis (Benjamini and Hochberg 1995).

To determine whether ASE measurements between genes within predicted operons were more similar those outside of operons, we performed 10,000 random samples of two categories of pairs of adjacent genes: either within predicted operons or outside of any predicted operon.

For each sampled pair of genes we calculated the difference in the absolute values of log_2_(ASE ratios). Finally, we asked whether the distribution of these differences from genes sampled within operons was significantly lower than that sampled outside of operons.

All statistical analyses were performed using R version 3.1.3 (R Core Team 2015). Kruskal-Wallis tests were performed using 10,000 permutations of the data as implemented in the ‘coin’ package (Hothorn et al. 2006).

### Detecting selection on cis-regulatory divergence

Gene Ontology (GO) categories for *H. volcanii* genes were obtained from the EBI Quick-GO database (accessed on 18 Feb. 2014; Binns et al. 2009). In the case of multi-gene operons, the operon was annotated as the union of the GO terms associated with its respective genes. For the purpose of interspecific comparisons, *H. mediterranei* orthologs were assigned to the same GO categories as *H. volcanii*.

Orthologs with significant *cis*-regulatory divergence at either level were divided into two categories based on the upregulating parental allele and ranked based on the magnitude of their absolute *cis* ratio (from largest to smallest). We searched for lineage-specific bias among GO biological process, GO molecular function, and GO cellular component. In order to detect lineage-specific bias within a gene set, we identified all functional categories containing at least 10 members in the set and determined whether significant bias existed in the direction of one or the other lineage using a χ^2^ ‘goodness of fit’ test. Because many different categories were being tested, we determined the probability of observing a particular enrichment by permuting ortholog assignments and repeating the test 10,000 times, retaining the most significant p-value observed in each functional dataset. We obtained a permutation-based p-value by asking how often a χ^2^ value of equal or greater significance would be observed in the permuted data. The test of bias was performed at two thresholds, using either the top 50% most biased orthologs, or analyzing all biased orthologs.

### Growth in low phosphate

The low phosphate media was Hv-Min medium (Allers et al. 2004), supplemented with potassium phosphate buffer (pH 7.5), the only phosphate source, to a final concentration of 0.1 mM phosphate. To compare the growth rates each strain was grown in low phosphate minimal broth medium at 42°C in shaking incubator for three days to reach OD_600_>0.4, then both strains diluted to be at the same OD (<0.15) to start the growth analysis. The growth curves were done using a Biotek ELX808IU-PC in 96-well plates at 42°C with continuous shaking, measuring OD_595_ every 30 minutes for 30 hours.

## Acknowledgements

We would like to thank the Fraser Lab for helpful discussions and advice, and R. Schreiber and S. Robinzon for technical assistance. This work was supported by NIH grant 1R01GM097171-01A1 and ISF grant 535/15.

---

**Supplemental Figure 1.**
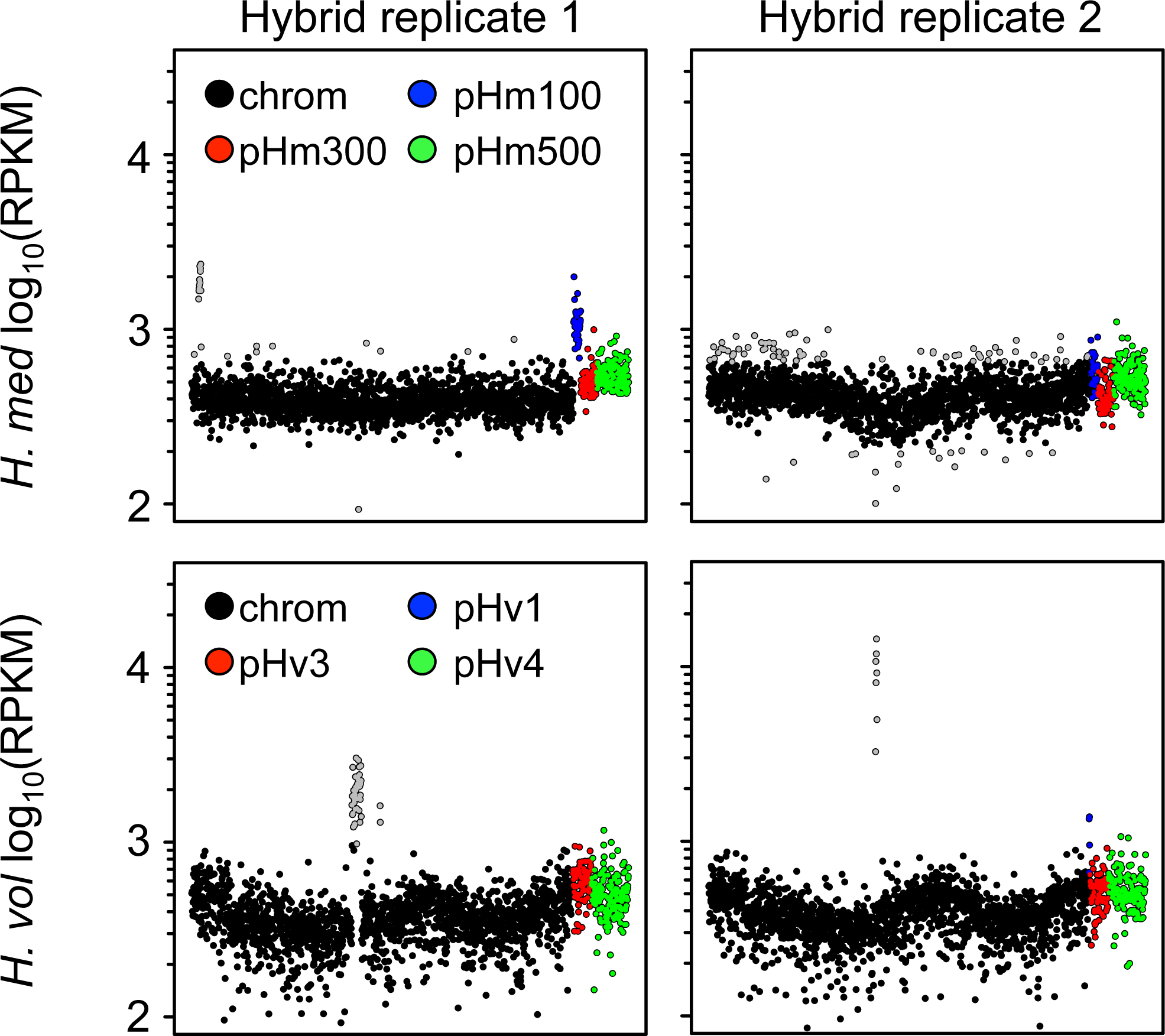
DNA-seq data reveals gene amplifications in both parental chromosomes, as well as hybrid specific variation in plasmid copy number. The *H. volcanii* chromosomes of both hybrid cultures as well as the *H. mediterranei* chromosome of replicate 1 revealed segmental amplifications spanning one or multiple genes (indicated in grey; see supplementary methods). Furthermore, the ratio of copy number among plasmids and main chromosomes varied between the two hybrids. Note that two-fold copy number variation is expected along chromosomes and plasmids as cultures were collected during active DNA replication during log-phase growth (Hawkins et al. 2013). med, mediterranei; vol, volcanii.

**Supplemental Figure 2.**
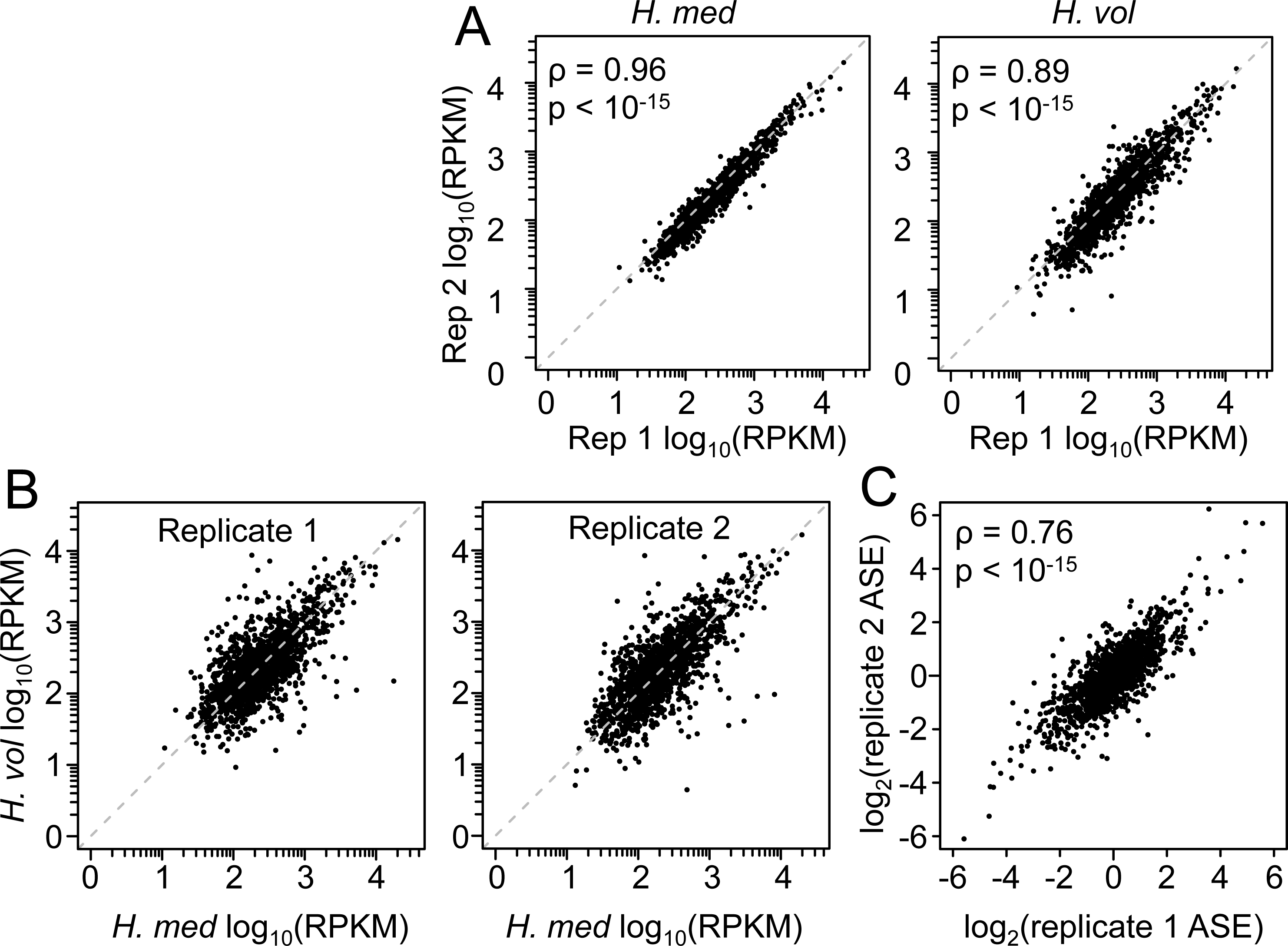
ASE measurements are concordant between biological replicates. Shown are the (A) ASE values comparing each species’ alleles across replicates; (B) ASE values comparing orthologs in each of the two biological replicates; and (C) as well as the inter-replicate correlation in ASE log-ratios. med, mediterranei; vol, volcanii.

**Supplemental Figure S3.**
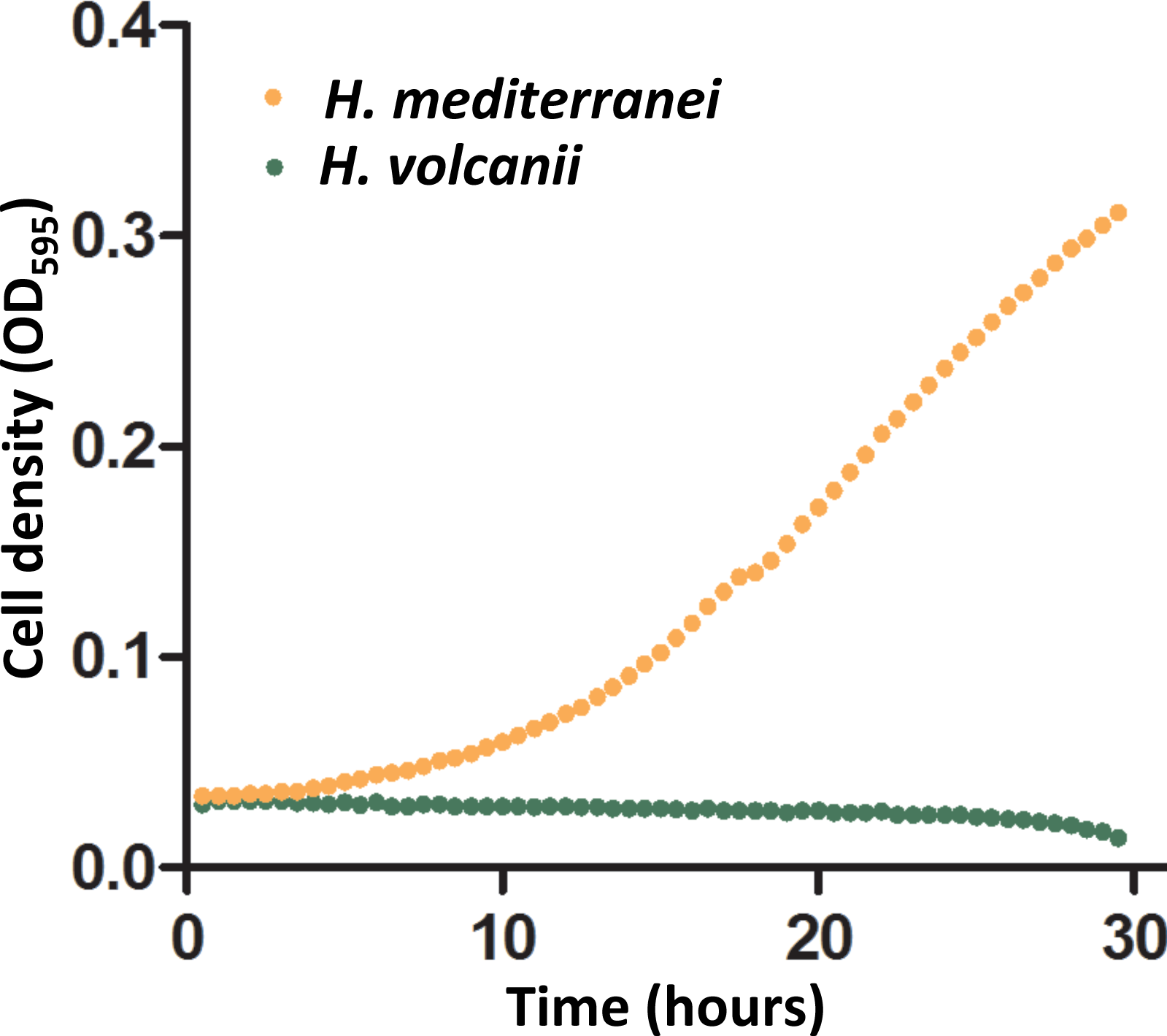
Biological replicate of growth curves in 0.1 mM phosphate.

